# Fast maturation of splenic dendritic cells upon TBI is associated with FLT3/FLT3L signaling

**DOI:** 10.1101/2021.11.29.470328

**Authors:** Jin Zhang, Zhenghui Li, Akila Chandrasekar, Shun Li, Albert Ludolph, Tobias Boeckers, Markus Huber-Lang, Francesco Roselli, Florian olde Heuvel

## Abstract

Systemic inflammatory consequences remain a significant burden after traumatic brain injury (TBI), with almost all organs affected. The spleen is connected with the brain by autonomic innervation and by soluble mediators, and the cross-talk between brain and spleen may be important to establish the systemic inflammatory response to TBI. Ethanol intoxication, the most common comorbidity of TBI, is posited to influence the peripheral inflammatory response either directly or through the brain-spleen cross-talk. Here we show that TBI causes a substantial change in transcription of genes associated with dendritic cells activation in the spleen, in particular a *FLT3*/*FLT3L* induction 3h after TBI, which was enhanced by EI. The *FLT3L* induction was associated with the phosphorylation of FLT3 receptor in CD11c+ dendritic cells, which enhanced the protein synthesis of a subset of mRNAs, as shown by the increase in pS6, peIF2A levels in dendritic cells. This corresponded to the upregulation of proteins associated with maturation process and immunostimulatory properties such MHC-II, LAMP1 and CD68, and of pro-inflammatory cytokines such as TNFα. Notably, EI enhanced the maturation of dendritic cells. However, whereas TBI decreases expression of the adrenergic 2b receptors on dendritic cells, EI increased it, thus augmenting the chances of cross-talk regulation of immune function by the autonomic system. In conclusion, this data indicates that TBI induces a fast maturation of the immunomodulatory functions of dendritic cells which is associated by FLT3/FLT3L signaling and which is enhanced by EI prior to TBI.

## 1. Introduction

Traumatic injury to the brain has acute, large-scale systemic consequences [1] that affect almost all organs and may lead to a compromised function of the heart, lung, gastrointestinal tract, liver, kidney, bones, lymphoid organs, and others, without direct systemic injury or infection [2, 3]. The systemic response to TBI is characterized by inflammation and, at the same time, a net systemic immunosuppression [4–6]. Although brain injury results in a systemic increase of inflammatory mediators and cytokines in both patients [7–9] and rodent models of TBI [10–12], there remains ample of evidence pointing towards a systemic immunosuppression post TBI, with a decrease in immune cells in the periphery [13–18]. Generalized immunosuppression is highly relevant on clinical grounds, since it may contribute to an enhanced vulnerability to infections observed after severe tissue injury.

The spleen is one of the most important immune regulatory organs, involved not only in red blood cell clearance but also in the facilitation of interactions between antigen presenting cells (APC) and T and B lymphocytes [19]. Prior investigation of the brain-spleen axis haven revealed interaction and regulation of splenic responses initiated by the central nervous system by either circulating mediators whose receptors are located on APCs [20] or by autonomic nerve fibers associated with splenic immune cells [21]. In fact, there have been reports of sympathetic and parasympathetic fibers innervating dendritic cells (DCs) in the spleen [22]. Furthermore, vagus nerve stimulation can reduce macrophage-induced TNF alpha release in the spleen through a so-called cholinergic anti-inflammatory pathway [23]. However, it remains poorly understood whether the spleen is directly and rapidly involved in the early response to TBI?

Ethanol intoxication (EI) is a frequent comorbidity in brain injury, with almost 40% of patients showing a positive blood alcohol level (BAL) upon admission [24]. Notably, the largest majority of patients showing EI in the context of TBI are not chronic alcohol consumers but rather young, and often episodic weekend drinkers (the so-called “drink-and-drive” patients). Recent studies revealed that acute EI can have beneficial effects on the neuroimmunological response following experimental TBI [25–27], these findings have been supported by clinical evidence [28–30]. However, some clinical studies have reported the opposite [31, 32]. Animal models have demonstrated that high doses of EI, reduce spleen size [33], reduce pro-inflammatory cytokines IL-1β and IL-6, but increase anti-inflammatory cytokine IL-10 [34]. Furthermore, EI has been shown to suppress antigen presentation by DCs [35], reflecting the immunosuppressive effects of ethanol intoxication on the spleen.

Is acute EI posited to interfere or rather to amplify the TBI-induced systemic immunoregulation, and in particular, is EI modulating TBI-induced immune reactions in the spleen? Given the high prevalence of EI in TBI patients, this question has direct clinical relevance. In this study we investigated the effect of a single high dose ethanol exposure prior to experimental-TBI in adult mice with focus on the immediate immunological responses in the spleen.

## 2. Material and Methods

### 2.1 Animals, traumatic brain injury model and ethanol treatment

This study represents a post-hoc analysis of spleen samples obtained from previous studies [25,26,36,37]. Investigations with these samples have never been reported before and this study was undertaken in accordance with the 3R-principle, to reduce the number of mice in animal experimentation but increase the scientific output from animal sacrifice. These experiments have been approved by Ulm University animal experimentation oversight committee and by the Regierungspräsidium Tübingen (license number 1222). Male wild-type mice (B6-SJL) were bred locally under standard housing conditions (24°C, 60-8% humidity, 12/12 light/dark cycle, with ad libitum access to food and water). TBI was performed on wild-type (WT) male mice aged p60-90, in agreement with epidemiological data in human TBI [38–40]. Experimental TBI was performed as previously reported [25,26,36,37]. Briefly, mice were administered buprenorphin (0.1mg/kg by subcutaneous injection) and anesthetized with sevoflurane (2.5% in 97.5% O_2_), after which the mice were subjected to a closed head weight drop TBI model. Animals were positioned in the weight-drop apparatus, and the TBI was delivered by a weight of 333 g free falling from a height of 2cm, targeting the parietal bone [41]. Directly after the TBI, mice were administered 100% O_2_ and apnea time was monitored. Control mice (sham group) had the same treatment and procedures (analgesia, anesthesia, skin incision and handling), but without the trauma being administered. Ethanol treatment was performed as previously described [42, 43]. Briefly, 100% synthesis grade ethanol was diluted in 0.9% NaCl saline to a final dilution of 32% volume/volume (32 µl of 100% ethanol and 68 µl of saline). Mice (20-25 g) were administered a volume 400-500 µl of diluted ethanol (to obtain a concentration of 5 g/kg) by oral gavage 30 min before TBI. Four experimental groups were investigated; saline-administered, subjected to sham surgery (saline-sham, SS), saline-administered, subjected to TBI (saline-TBI, ST), ethanol administered, subjected to sham surgery (ethanol-sham, ES), ethanol administered, subjected to TBI (ethanol-TBI, ST).

### 2.2 Tissue isolation

Three hours post trauma, mice were euthanized by cervical dislocation and organs were harvested for further processing. The spleen was dissected quickly and snap frozen in dry ice for further processing. Tissue was used for either RNA isolation and Quantitative RT-PCR, for tissue sectioning and immunofluorescence staining.

### 2.3 RNA isolation and Quantitative RT-PCR

RNA was isolated from the spleen using QIAzol (Qiagen, Germany) by disrupting and homogenizing the tissue in 1mL QIAzol, after which 200µl of chloroform was added and vortexed for 15 sec. The samples were placed at RT for 10 min and centrifuged for 10 min 12.000g at 4⁰C to achieve the phase separation. The top layer (containing RNA) was moved to another tube and precipitated with the same amount of isopropanol. The samples were placed at RT for 10 min and centrifuged for 10 min 12.000g at 4⁰C. The isopropanol was removed and 1mL of 75% ethanol in DEPC treated dH_2_0 was added and mixed. The samples were centrifuged for 10 min 8000 x g at 4⁰C, ethanol was removed and the samples air dried. The RNA pellet was redissolved in 20µl RNAse-free dH_2_O. RNA concentration was determined by Nanodrop. Reverse transcription was performed by adding 5µl random hexamers (Biomers, Germany) to 0,75μg RNA (total volume 40µl diluted in dH_2_O) and incubated for 10 min at 70⁰C. The samples were placed on ice and a mastermix of 0.5µl reverse transcriptase (Promega, germany), 0.5µl RNase Inhibitor (RiboLock, Thermo Scientific, Germany), 2µl dNTPs (genaxxon, Germany) and 12µl reverse transcriptase buffer (Promega, Germany). The samples were incubated for 10 min at RT and placed in a water bath for 45 min at 42⁰C. The samples were incubated for 3 min at 99⁰C, placed on ice and frozen until further use. qPCR was performed on the Light Cycler 480II (Roche) with the Power PCR TB green PCR master mix (Takara, Japan). 2µl of sample cDNA was used in a total volume of 10µl (3µl primer mix and 5µl of TB green) in a 96-well plate, all samples were duplicated and the housekeeping gene GAPDH was used as a control (for a complete overview of cytokine sequences see Supplementary Table 1). The Ct values obtained from the lightcycler were normalized according to the following equation: 2-ΔCt (ΔCt = Cttarget gene -CtGapdh) = relative mRNA.

### 2.4 Tissue sectioning and immunofluorescent staining

Frozen spleen tissue was embedded in OCT (tissue tek, The Netherlands), 10 micron sections were cut with the cryostat and mounted on glass slides. Slides were stored for 24h at -80⁰C and washed in 1x PBS, followed by a 10 min fixation step of the sections in 4% PFA. Target retrieval was performed in sodium citrate buffer pH 8.5, followed by blocking of the sections in blocking buffer (3% BSA, 0.3% triton X-100; PBS) for 2h at RT. Primary antibodies (for a complete overview of antibodies used see Supplementary Table 2) were diluted in blocking buffer and incubated for 48h at 4⁰C, followed by 3 x 30min washes in PBS at RT. Secondary antibodies were diluted in blocking buffer and incubated for 2h at RT, followed by 3 x 30 min washes in PBS, the sections were mounted using Fluorogold prolong antifade mounting medium (Invitrogen, Germany).

### 2.5 single mRNA in situ hybridization

Fluorescent in situ mRNA hybridisation was used as previously reported [44] in agreement to the manufacturer’s instructions (ACDBio, RNAscope, Fluorescent In Situ Hybridisation for fresh frozen tissue, all reagents and buffers were provided by ACDBio), with small modifications [25]. Briefly, frozen spleen tissue was embedded in OCT (tissue tek, the Netherlands), 10 micron sections we cut with the cryostat and mounted on super frost plus glass slides. Slides were stored for 24h at -80⁰C and fixed for 10 min in 4% PFA at 4⁰C. Sections were covered by Protease IV and incubated for 30 min at 40⁰C followed by 2 x 2 min washing in PBS. After which the probe (TNFα and ARB2) has been added and incubated for 4.5h at 40⁰C followed by 2 x 2 min washing step with wash buffer. Then, amplification 1 was added to the sections and incubated for 30 min at 40⁰C followed by 2 x 2 min washing step with wash buffer. After which, amplification 2 was added to the sections and incubated for 15 min at 40⁰C followed by 2 x 2 min washing step with wash buffer. As a final amplification step, amplification 3 was added and incubated for 30 min at 40⁰C followed by 2 x 2 min washing step with wash buffer. Finally, the detection step was performed by adding detection reagent 4A to the sections and incubated for 45 min at 40⁰C followed by 2 x 2 min washing step with wash buffer. After which the sections were blocked in blocking buffer (3% BSA, 0.3% triton X-100; PBS) for 1h at RT followed by an overnight incubation with primary antibodies diluted in blocking buffer. The sections were washed 3 x 30 min in PBST and incubated with secondary antibodies diluted in blocking buffer for 2h at RT. A final washing step 3 x 30 min in PBST was performed and the sections were counterstained with DAPI and mounted using Fluorogold prolong antifade mounting medium (Invitrogen, Germany)

### 2.5 Image acquisition and image analysis

Immunofluorescent staining was imaged with a keyence BZ-X800 microscope (Keyence, Japan) equipped with a 100x oil objective, a single optical section with 3×3 tile scan was made spanning an area covering a splenic follicle and the red pulp. Acquisition parameters were set to avoid hyper- or hypo-saturation and kept constant for each experimental set. Images were merged with the BZ-II analyzer software (Keyence, Japan) and analyzed using the Image-J. Fluorescent intensity was assessed by manually tracing the CD11c+ cells and measuring the mean gray value. Density was assessed by using the Image-J plugin cell counter. The analyzed marker was assessed as high or low expressing, by thresholding the signal. Each picture was analyzed by making a ratio between total CD11c+/CD45+ cells and the imaged marker or the ratio between CD11c-/CD45+ cells and the imaged marker.

Single mRNA in situ Hybridization images were acquired using an LSM-710 (Carl Zeiss, Germany) microscope with an 40x oil objective with optical thickness fitted to the optimum value. A z-stack of 8 images has been acquired at 1024×1024 pixel resolution and 16-bit depth. Acquisition parameters were set to avoid over and under-saturation and kept the same for each experimental set. TNFα and ADRB2 mRNA density in CD11c+ cells were assessed by using the Image-J plugin cell counter.

### 2.6 Statistical analysis

Statistical analysis for the gene expression data sets (Fig. 1 and Fig 2.) was performed using the IBM software suite, using a Two-way Multivariate ANOVA (with Wilk’s λ parameter), because the experiment included multiple dependent variables (cytokines or chemokines) and two independent variables (TBI and ethanol treatment). The post-hoc comparisons were performed with Two-way ANOVA with Tukey post-hoc correction. All groups were tested for normality using the Shapiro-Wilk test. For the histological datasets, Two-way ANOVA was performed with Tukey post-hoc comparison, since two independent treatments were done (saline/ethanol and sham/TBI). Statistical significance was set at p < 0.05 after multiple-comparison correction.

**Figure 1:**
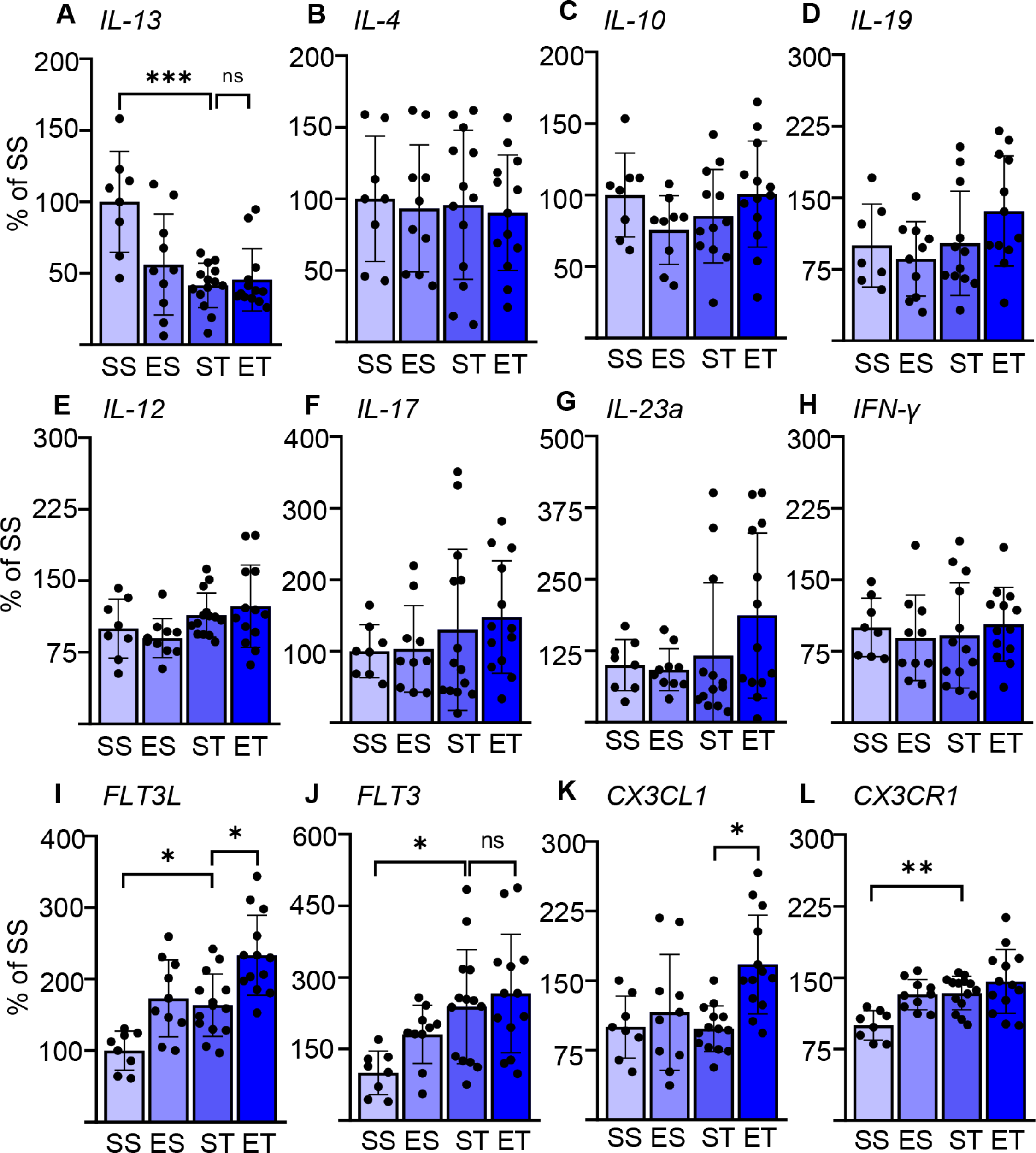
EI enhances the selective cytokine expression post TBI. Cytokine expression screening in the spleen of saline sham (SS), ethanol sham (ES), saline TBI (ST) and ethanol TBI (ET) treated mice 3h after trauma. A-D) Barplots show relative expression of Th2-cell, anti-inflammatory markers; *IL-13*, *IL-4*, *IL-10* and *IL-19*. *IL-13* expression showed a significant downregulation after TBI (SS vs ST; P < 0.0005), ethanol pretreatment did not alter the TBI induced effect on *IL-13* (ST vs ET; p > 0.05). *IL-4*, *IL-10* and *IL-19* were not affected by any treatment. E-H) Barplots show relative expression of Th1-cell, pro-inflammatory markers; *IL-12*, *IL-17*, *IL-23a* and *IFN-y*. TBI with or without ethanol pretreatment showed no significant differences. I-L) Barplots show relative expression of DCs-monocyte specific mediators; *FLT3*, *FLT3R*, *CX3CL1*, *CX3CR1*. TBI resulted in a significant increase of *FLT3* (SS vs ST; p < 0.037), *FLT3L* (SS vs ST; p < 0.045) and *CX3CR1* (SS vs ST; p < 0.01). Ethanol pretreatment resulted in a further significant enhancement of *FLT3* (ST vs ET; p < 0.026) and *CX3CL1* (ST vs ET; p < 0.006). Data shown as barplots and individual data points. Group size: SS N = 8, ES N = 10, ST N = 14, ET N = 13. *: p < 0.05; **: p < 0.01; ***: p < 0.001.

**Figure 2:**
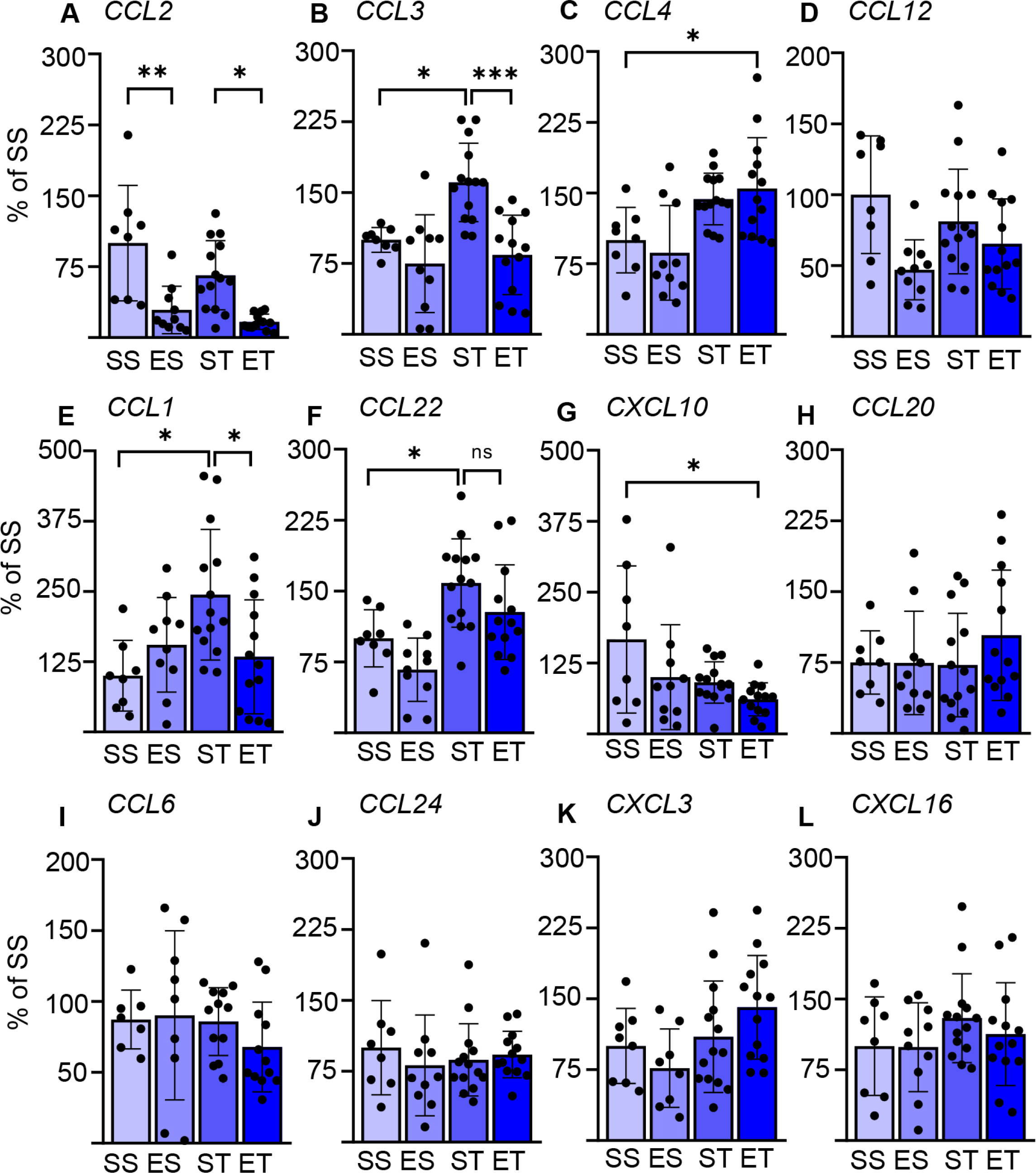
Splenic chemokine expression shows a fast response to TBI and EI. Chemokine expression data in the spleen of saline sham (SS), ethanol sham (ES), saline TBI (ST) and ethanol TBI (ET) treated mice 3h after trauma. A-D) Barplots show relative expression of modulators of DC biology; *CCL2*, *CCL3*, *CCL4* and *CCL12*. TBI resulted in a significant upregulation of *CCL3* (SS vs ST; p = 0.028), ethanol pretreatment significantly reduced the TBI induced *CCL3* expression (ST vs ET; p = 0.0006). Ethanol by itself resulted in a significant downregulation of *CCL2*, independent of TBI (SS vs ES; p = 0.0021; ST vs ET; p = 0.014). EI together with TBI resulted in a significant upregulation of *CCL4* (SS vs ET; p = 0.042). *CCL12* expression was not affected by any treatment. E-H) Barplots show relative expression of modulators of Th-cells; *CCL1*, *CCL22*, *CXCL10* and *CCL20*. TBI resulted in a significant upregulation of *CCL1* (SS vs ST; p = 0.019) and *CCL22* (SS vs ST; p = 0.039), ethanol pretreatment significantly reduced the TBI induced *CCL1* expression (ST vs ET; p = 0.037) but not for *CCL22* (ST vs ET; p = 0.35). EI together with TBI resulted in a significant downregulation of *CXCL19* (SS vs ET; p = 0.014). *CCL10* expression was not affected by any treatment. I-L) Barplots show relative expression of modulators of leukocytes and NK-cells; *CCL6*, *CCL24*, *CXCL3* and *CXCL16*. TBI with or without ethanol pretreatment showed no significant differences. Data shown as barplots and individual data points. Group size: SS N = 8, ES N = 10, ST N = 14, ET N = 13. *: p < 0.05; **: p < 0.01; ***: p < 0.001.

## 3. Results

### 3.1 TBI induces a rapid and selective induction of cytokine upregulation in the spleen, enhanced by concomitant EI

Systemic immune functions are quickly modulated by the occurrence of neurological conditions through pathways conceptualized as the “brain-spleen axis” [45–47]. We set out to verify if any rapid modification in the splenic immune responses would take place upon mild/moderate TBI and, most importantly, if the comorbidity of EI would significantly interact with such responses.

As an entry point, we screened the induction of the mRNA of several cytokines in whole-spleen extracts obtained 3h after TBI (or sham surgery), pretreated (30 min) either with saline or with ethanol (5g/kg).

We considered three sets of cytokines: prototypical Th1-cellular-immunity-directed, pro-inflammatory mediators (*IL-12*, *IL-17*, *IL-23a* and *IFN-y*), prototypical Th2-B-cells-directed, anti-inflammatory mediators (*IL-4*, *IL-10*, *IL-13* and *IL-19*) and prototypical DCs-monocytes mediators (*FLT3*, *FLT3L*, *CX3CL1* and *CX3CR1* [48–51]. Surprisingly, we identified a significant effect of both the TBI itself and EI (Two-way MANOVA; TBI: F = 7.379; Wilks λ = 0.253; p = 0.0001; EI: F = 4.599; Wilks λ = 0.352; p = 0.0001), the interaction of the two parameters showed no significance in the Two-way MANOVA (F = 1.801; Wilks λ = 0.581; p = 0.09). This could be attributed (Two-way ANOVA, Tukey corrected) to the significantly increased expression of *FLT3L* upon TBI alone (F_(3, 34)_ = 9.771; SS vs ST; 100±27 vs 163±44; p = 0.045; Fig. 1I), *FLT3* (F_(3, 36)_ = 4.266; SS vs ST; 100±46 vs 239±119; p = 0.037; Fig. 1J) and *CX3CR1* (F_(3,41)_ = 6.825; SS vs ST; 100±16 vs 134±18; p = 0.01; Fig. 1L), whereas *IL-13* was significantly downregulated (F_(3, 37)_ = 7.499; SS vs ST; 100±35 vs 42±16; p = 0.0005; Fig. 1A). Ethanol intoxication before TBI resulted in a significant further enhancement of the expression of *FLT3L* (ST vs ET; 163±44 vs 233±56; p = 0.026; Fig. 1I) and *CX3CL1* (ST vs ET; 98±25 vs 168±53; p = 0.006; Fig. 1K), whereas *FLT3* and *CX3CR1* were unaltered in EI-TBI compared to saline-TBI (*FLT3*: ST vs ET; 239±119 vs 266±124; p = 0.93; Fig. 1J; *CX3CR1*: ST vs ET; 134±18 vs 146±34; p = 0.52; Fig. 1L). Most notably, no effect was observed on any of the other cytokines considered.

These screening data not only point towards a rapid activation of splenic immune cells upon TBI, but also reveal a pattern compatible with a selective effect on innate immunity.

### 3.2 Rapid modulation of the splenic chemokine pattern by TBI and by EI/TBI

We sought to confirm and further extend the rapid effect of TBI on splenic immune cells, in particular on DCs. We assessed the expression of a set of chemokines known to be strong modulators of innate immune responses [52, 53]. The focus was set, in particular on DC biology (and other APC: *CCl2*, *CCL3*, *CCl4* and *CCL12*; [54–56]) Th cells (*CCL1*, *CCL20*, *CCL22* and *CXCL10*; [57–60]) or leukocytes and NK cells (*CCL6*, *CCl24*, *CXCL3* and *CXCL16*; [61–64]). We could identify a significant effect of TBI (Two-way MANOVA: F = 4.494; Wilks λ = 0.357; p = 0.0001) and EI (F = 4.956; Wilks λ = 0.335; p = 0.0001), the interaction of the two parameters showed no significant effect in the Two-way MANOVA (F = 1.776; Wilks λ = 0.585; p = 0.099). However, post-hoc analysis (Tukey corrected) showed that TBI alone significantly increased the expression of *CCL3* (F_(3, 37)_ = 8.910; SS vs ST; 100±13 vs 161±42; p = 0.028; Fig. 2B), *CCL1* (F_(3, 39)_ = 4.156; SS vs ST; 100±63 vs 244±116; p = 0.019; Fig. 2E), and *CCL22* (F_(3, 37)_ = 7.483; SS vs ST; 100±30 vs 159±47; p = 0.039; Fig. 2F). On the other hand, concomitant EI significantly downregulated *CCL3* (ST vs ET; 161±42 vs 84±42; p = 0.0006; Fig. 2B) and *CCL1* (ST vs ET; 244±116 vs 134±101; p = 0.037; Fig. 2E), but *CCL22* was unaffected by EI and remained upregulated (ST vs ET; 159±47 vs 128±50; p = 0.35; Fig. 2F). The EI/TBI group presented a distinct chemokine profile: a significant upregulation of *CCL4* was observed only in the EI/TBI group but not in the TBI and EI groups alone ((F_(3, 39)_ = 6.093; SS vs ET; 100±35 vs 155±54; p = 0.042; Fig. 2C). Conversely, the EI/TBI group displayed the downregulation of *CXCL10* expression (F_(3, 41)_ = 3.458; SS vs ET; 100±78 vs 37±18; p = 0.014; Fig. 2G). Finally, ethanol exposure alone caused a downregulation of *CCL2* in both sham and TBI groups (F_(3, 36)_ = 9.679; SS vs ES; 100±61 vs 29±25; p = 0.0021; ST vs ET; 66±37 vs 17±8; p = 0.014; Fig. 2A).

These data not only confirm the rapid engagement of the brain-spleen axis upon TBI but also show that EI displays a substantial modulatory effect on innate and adaptive immunity, in particular on APCs.

### 3.3 TBI induces the phosphorylation of FLT3 and its downstream signaling partner BTK in DCs, which are both enhanced by EI

Our expression screening indicated a possible upregulation of FLT3 signaling in association with a cytokine pattern compatible with the involvement of DCs. Considering the well-known role of FLT3/FLT3L in regulating DC immunobiology [65], we sought direct confirmation of enhanced Flt3 engagement in splenic DCs upon TBI and EI/TBI. For this aim, we immunostained thin sections of the spleen from the four treatment groups for the pan-DCs marker CD11c [21], for the pan-leukocyte marker CD45 [66] and for phosphorylated FLT3 (pFLT3, Y589/591; Fig. 3A). Phospho-FLT3 immunoreactivity was highly inhomogeneous in the spleen sections, with a comparatively small number of cells highly positive for pFLT3 localized around follicles and a minor number of cells with moderate pFLT3 immunoreactivity scattered in the parenchyma. Initial immunostainings revealed that CD11c density was not altered by TBI or concomitant EI (Two-way ANOVA, F_(3, 12)_ = 0.9360; p = 0.45; Supplementary Fig. 1B-C). Co-immunostaining with CD11c and CD45 revealed that nearly all the pFLT3^high^ cells (upon binning) were CD45+ (supplementary Fig. 1D-E). However, when investigating the density of pFLT3^high^ cells in CD45+/CD11c- cells, we found no significant difference among treatment groups (Two-way ANOVA, F_(3, 12)_ = 0.8446; p = 0.50; Supplementary Fig. 1D-E). Although the fluorescent intensity of pFLT3 in CD11c+ DCs was not altered across the four treatment groups (Two-way ANOVA, F_(3, 588)_ = 2.541; p = 0.06; Fig. 3B), the number of pFLT3^high^ /CD11c+ cells was significantly affected by TBI and treatment (Two-way ANOVA, F_(3, 12)_ = 45.84; p < 0.0001) and post-hoc comparison (Tukey corrected) revealed a significant increase in pFLT3^high^/CD11c+ cells after TBI alone (SS vs ST; 20±1% vs 48±4%; p < 0.0001; Fig. 3C) and, interestingly, a substantial further enhancement in EI-TBI (ST vs ET; 48±4% vs 65±3%; p = 0.0061; Fig. 3C), confirming that TBI alone and in particular EI-TBI strongly induces FLT3 signaling in DCs. In order to verify that the phosphorylation of the FLT3 receptor corresponded to the functional engagement of signal transduction pathways, we monitored the phosphorylation of the Bruton’s tyrosine kinase (BTK), an established downstream target of FLT3 [67]. Immunostaining of spleen thin sections for phosphorylated BTK (pBTK, Y223; Fig. 3D) revealed a pattern closely resembling that of pFLT3. Fluorescent intensity of pBTK in CD11c+ cells showed a significant difference between treatment groups (Two-way ANOVA, F_(3, 444)_ = 123.5; p < 0.0001) and a post-hoc comparison (Tukey corrected) showed a significant increase after TBI (SS vs ST; 44±9 vs 48±9; p = 0.0008; Fig. 3E). EI resulted in a further increase of pBTK fluorescent intensity (ST vs ET; 48±9 vs 62±11; p < 0.0001; Fig. 3E). Likewise, quantification of the number of pBTK^high^/CD11c+ cells revealed a significant effect in the treatment groups (Two-way ANOVA, F_(3, 12)_ = 68.07; p < 0.0001), due to the massive increase of pBTK^high^/CD11c+ cells in the Saline-TBI compared to the sham group (Tukey’s post-hoc; SS vs ST; 31±6% vs 58±5%; p < 0.0001; Fig. 3F), which was further increased in the EI/TBI group (ST vs ET; 58±5% vs 67±2%; p = 0.022; Fig. 3F). Taken together, these data demonstrate that TBI causes fast (3h) elevation of the phosphorylation of the FLT3 receptor (and its downstream target BTK) in splenic DCs, and that this effect is substantially amplified by concomitant EI.

**Figure 3:**
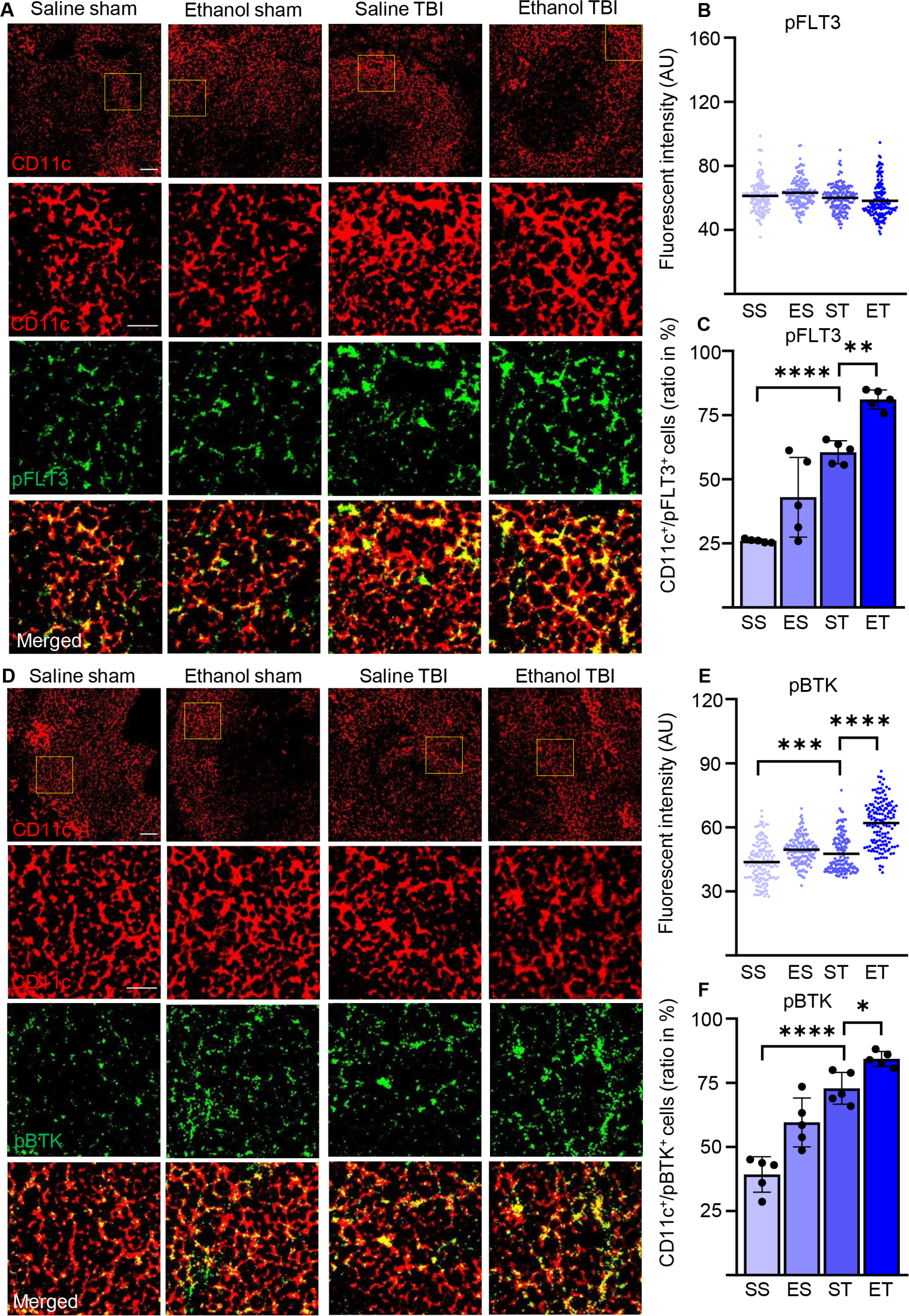
TBI induced phosphorylation of FLT3 and downstream signaling partner BTK is further enhanced by EI. Immunofluorescence staining of pFLT3 and pBTK with DC marker CD11c on thin spleen sections of saline sham (SS), ethanol sham (ES), saline TBI (ST) and ethanol TBI (ET) treated mice 3h after trauma. A-C) immunofluorescent staining of pFLT3 colocalized with CD11c resulted in no significant difference in fluorescent intensity between treatment groups (p = 0.06). However, the amount of FLT3^high^/CD11c+ cells revealed a significant increase after TBI (SS vs ST; p < 0.0001) and a further significant enhancement upon EI (ST vs ET; p = 0.006). D-F) immunofluorescent staining of pBTK colocalized with CD11c resulted in a significant increase in fluorescent intensity upon TBI (SS vs ST; p = 0.0008), with a further significant increase after EI (ST vs ET; p = 0.006). Likewise, the number of BTK^high^/CD11c+ cells revealed a significant increase after TBI (SS vs ST; p < 0.0001) with a further significant increase after EI (ST vs ET; p < 0.0001). Data shown as scatterplots or barplots with individual data points. Group size: SS N = 5, ES N = 5, ST N = 5, ET N = 5. *: p < 0.05; **: p < 0.01; ***: p < 0.001; ****: p < 0.0001. Scale bar overview: 50 µm; scale bar insert: 20 µm.

### 3.4 TBI and concomitant EI-TBI strongly induce protein synthesis in splenic DCs

Activation of DCs is characterized by a substantial remodelling of their metabolic rates and is in particular associated with the upregulation of protein synthesis [68] which, in turn, brings about the long-term adaptation of the cellular metabolism to the immune function [69]. Since FLT3 is a strong regulator of mTor [70], a major driver of protein synthesis, we set out to determine if the upregulation of FLT3 signaling observed upon TBI, and enhanced by EI/TBI, was accompanied by the activation-associated increase in protein synthesis. First, we considered the levels of phosphorylated S6 ribosomal protein (pS6), a proxy of mTOR activation directly involved in increasing ribosomal translation of mRNA. Immunostaining of thin spleen sections for phosphorylated-S6 (pS6, S235/236; Fig. 4A) revealed a diffuse immunopositivity in the cytoplasm of close to every cell; however, upon TBI, CD11c+ cells stood out for having a massive increase in pS6 immunofluorescence intensity (Two-way ANOVA, F_(3, 465)_ = 418.3; p < 0.0001). The post-hoc comparison (Tukey corrected) showed a significant increase after TBI (SS vs ST; 44±6 vs 70±13; p < 0.0001; Fig. 4B), further enhanced by EI (ST vs ET; 70±13 vs 98±23; p < 0.0001; Fig. 4B). This effect corresponded to the significant increase in the number of CD11c+ displaying high levels of pS6 (Two-way ANOVA, F_(3, 12)_ = 26.09; p < 0.0001) due to the substantial elevation of pS6 occurring after TBI (Tukey’s post-hoc; SS vs ST; 32±4% vs 58±8%; p = 0.0002; Fig. 4C) but not further increased by EI/TBI compared to TBI alone (ST vs ET; 58±8% vs 65±7%; p = 0.34; Fig. 4C). Thus, EI magnifies the increased pS6 levels in all sensitive CD11c+ cells upon TBI but does not increase the number of cells responding to TBI itself.

**Figure 4:**
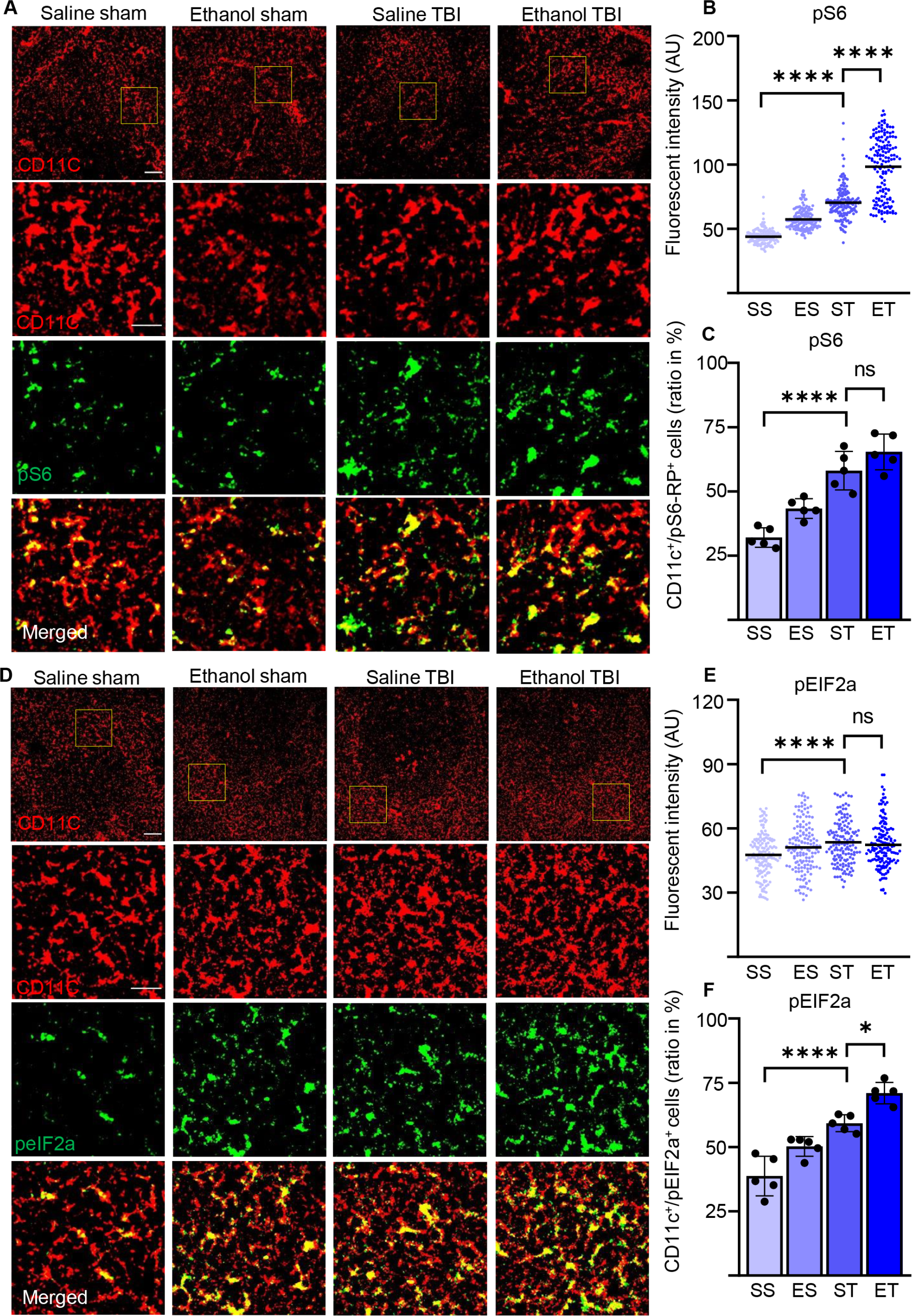
TBI induced metabolic rate and protein synthesis is further enhanced by EI. Immunofluorescence staining of pS6-RP and peIF2A with DC marker CD11c on thin spleen sections of saline sham (SS), ethanol sham (ES), saline TBI (ST) and ethanol TBI (ET) treated mice 3h after trauma. A-C) immunofluorescent staining of pS6-RP colocalized with CD11c resulted in a significant increase in fluorescent intensity upon TBI (SS vs ST; p < 0.0001), with a significant enhancement after EI (ST vs ET; p < 0.0001). The amount of pS6-RP^high^/CD11c+ cells revealed a significant increase after TBI (SS vs ST; p < 0.0001) however, no significant difference upon EI (ST vs ET; p = 0.34). D-F) immunofluorescent staining of peIF2A colocalized with CD11c resulted in a significant increase in fluorescent intensity upon TBI (SS vs ST; p < 0.0001), with no difference after EI (ST vs ET; p = 0.78). The number of peIF2A^high^/CD11c+ cells revealed a significant increase after TBI (SS vs ST; p = 0.0001) with a further significant increase after EI (ST vs ET; p = 0.0107). Data shown as scatterplots or barplots with individual data points. Group size: SS N = 5, ES N = 5, ST N = 5, ET N = 5. *: p < 0.05; **: p < 0.01; ***: p < 0.001; ****: p < 0.0001. Scale bar overview: 50 µm; scale bar insert: 20 µm.

Furthermore, the levels of the cellular stress-related phospho-eIF2A [68, 71] show a significant difference in fluorescence intensity in CD11c+ cells within treatment groups (Two-way ANOVA, F_(3, 444)_ = 8.553; p < 0.0001) due to a significant increase upon TBI (Tukey’s post-hoc; SS vs ST; 48±10 vs 54±10; p < 0.0001; Fig. 4E), again with no difference between EI/TBI and TBI (ST vs ET; 54±10 vs 52±11; p = 0.78; Fig. 4E). Colocalization of peIF2A^high^ cells with CD11c+ cells reveal a significant difference within treatment groups (Two-way ANOVA, F_(3, 12)_ = 40.50; p < 0.0001). The post-hoc comparison (Tukey corrected) shows a significant increase after TBI (SS vs ST; 39±8% vs 60±4%; p = 0.0001; Fig. 4F) with a further increase in EI/TBI compared to TBI alone (ST vs ET; 60±4% vs 67±6%; p = 0.01; Fig. 4F). Thus, TBI induces a substantial increase in protein synthesis, with some alteration of cap-dependent translation in DCs, and this effect is amplified by EI.

### 3.5 TBI and EI cooperate to enhance the maturation of splenic DCs

DC maturation occurs on stimulation with pathogens associated molecular patterns or danger associated molecular patterns (PAMPs or DAMPs; [72, 73]). Maturation of DCs is accompanied by an increase of antigen presentation and by lysosomal activity. Antigen presentation of DCs (and macrophages) can be visualized by investigating MHC-complexes in CD11c+ cells [74]. We wondered if the signaling events and metabolic reprogramming observed in splenic DCs after TBI also corresponded to the appearance of a “mature” APC phenotype. Spleen sections were stained for MHC-II and CD11c (Fig. 5A), which reveals a high amount of MHC-II in DCs upon TBI, resulting in a significant increase of fluorescent intensity within treatment groups (Two-way ANOVA, F_(3, 444)_ = 118.0; p < 0.0001). Post-hoc analysis (Tukey corrected) indicated a significant increase after TBI (SS vs ST; 26±9 vs 41±11; p < 0.0001; Fig. 5B) with a further increase after EI (ST vs ET; 41±11 vs 45±13; p < 0.0001; Fig. 5B). Furthermore, when assessing the density of colocalizing MHC-II^high^ with CD11c+ cells, we found a significant increase within treatment groups (Two-way ANOVA, F_(3, 12)_ = 60.83; p < 0.0001) due to a massive increase upon TBI (Tukey corrected; SS vs ST; 44±6% vs 74±5%; p < 0.0001; Fig. 5C) and a further increase after EI (ST vs ET; 74±5% vs 84±4%; p = 0.045; Fig. 5C). Thus, both the number of CD11c+ cells expressing MHC-II and the levels of such expression increase upon TBI and are further increased by EI.

**Figure 5:**
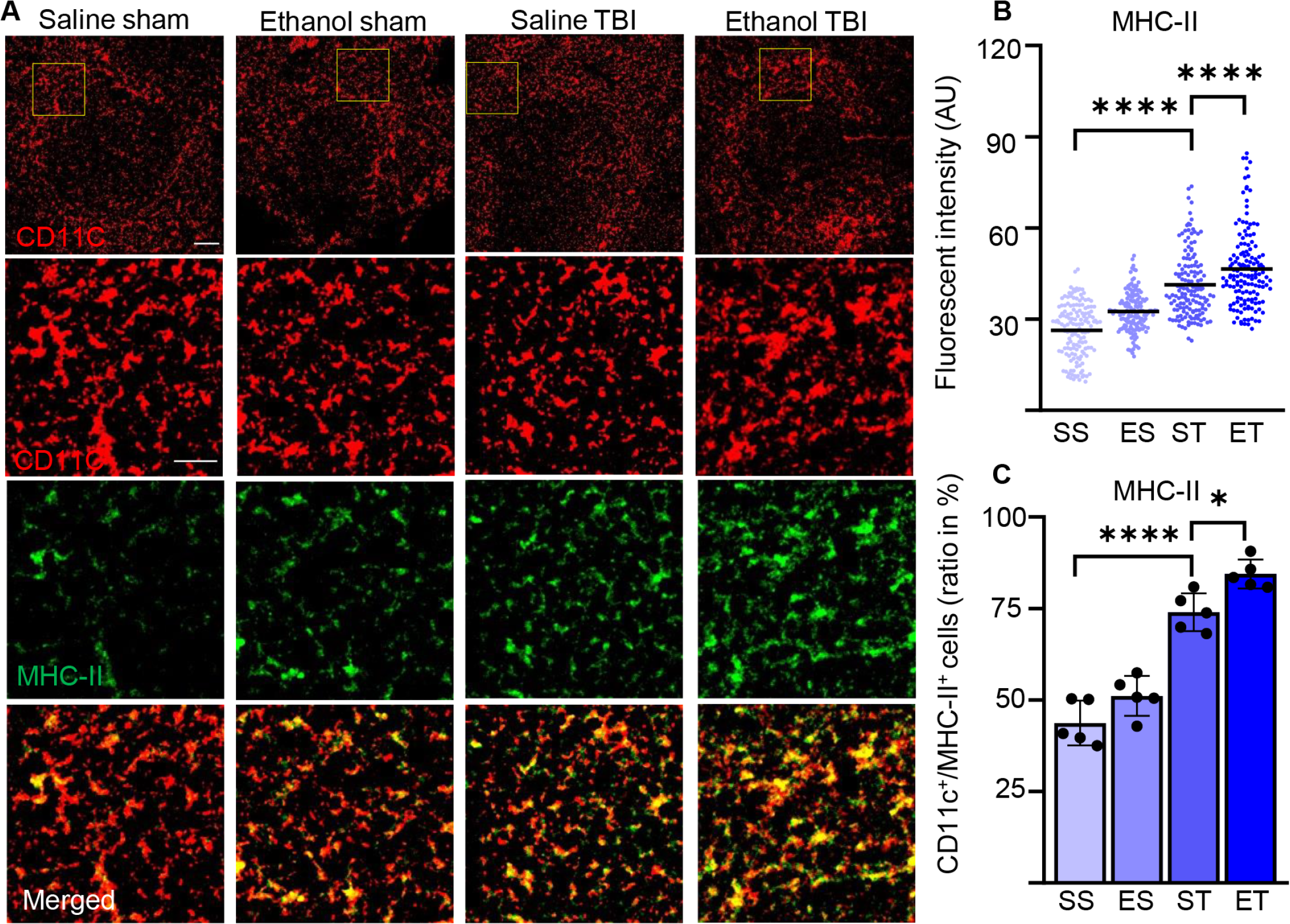
TBI induces antigen presentation on DCs, which is further enhanced by EI. Immunofluorescence staining of MHC-II with DC marker CD11c on thin spleen sections of saline sham (SS), ethanol sham (ES), saline TBI (ST) and ethanol TBI (ET) treated mice 3h after trauma. A-C) immunofluorescent staining of MHC-II colocalized with CD11c resulted in a significant increase in fluorescent intensity upon TBI (SS vs ST; p < 0.0001), with a significant enhancement after EI (ST vs ET; p < 0.0001). Likewise, the amount of MHC-II^high^/CD11c+ cells revealed a significant increase after TBI (SS vs ST; p < 0.0001) with a significant enhancement upon EI (ST vs ET; p = 0.045). Data shown as scatterplots or barplots with individual data points. Group size: SS N = 5, ES N = 5, ST N = 5, ET N = 5. *: p < 0.05; **: p < 0.01; ***: p < 0.001; ****: p < 0.0001. Scale bar overview: 50 µm; scale bar insert: 20 µm.

We also assessed the lysosomal activity of splenic DCs by investigating CD68 and LAMP1 in our samples [75, 76]. Immunostaining of CD68 and CD11c (Fig. 6A) on spleen sections revealed a significant difference in intensity within treatment groups (Two-way ANOVA, F_(3, 444)_ = 79.68; p < 0.0001) with a strong increase after TBI (Tukey corrected, SS vs ST; 36±13 vs 51±13; p < 0.0001; Fig. 6B) and an even further enhancement upon EI in TBI (ST vs ET; 51±13 vs 63±22; p < 0.0001; Fig. 6B). When assessing the density of CD68^high^ cells colocalized with CD11c+ DCs, the treatment resulted in a significant difference within groups (Two-way ANOVA, F_(3, 12)_ = 41.25; p < 0.0001) due to TBI which resulted in an increase in the amount of CD68^high^/CD11c+ cells in the spleen (SS vs ST; 25±9% vs 36±6%; p = 0.0007; Fig. 6C) and a further significant increase in EI (ST vs ET; 36±6% vs 46±9%; p = 0.002; Fig. 6C). Similarly, immunostaining of LAMP1 and CD11c (Fig. 6D) on spleen sections revealed a significant effect in intensity within treatment groups (Two-way ANOVA, F_(3, 444)_ = 214.9; p < 0.0001) due to a significant increase upon TBI (Tukey corrected; SS v ST; 24±6 vs 38±12; p < 0.0001; Fig. 6E) and a further significant increase was detected after EI (ST vs ET; 38±12.3 vs 49.2±7.6; p < 0.0001; Fig. 6E). The density of LAMP1^high^ cells colocalized with CD11c+ cells exhibited a significant effect within the treatment groups (Two-way ANOVA, F_(3, 12)_ = 43.02; p < 0.0001). The post-hoc comparison within the treatment groups (Tukey corrected) showed a strong significant upregulation after TBI (SS vs ST; 30±4% vs 54±6%; p = 0.0007; Fig. 6F) and a further significant enhancement due to EI (ST vs ET; 54±6% vs 67±8%; p = 0.01; Fig. 6F). The convergence of MHC-II, LAMP-1 and CD68 upregulation indicate that upon TBI, APC functions are upregulated in splenic DCs and they are further enhanced by concomitant EI.

**Figure 6:**
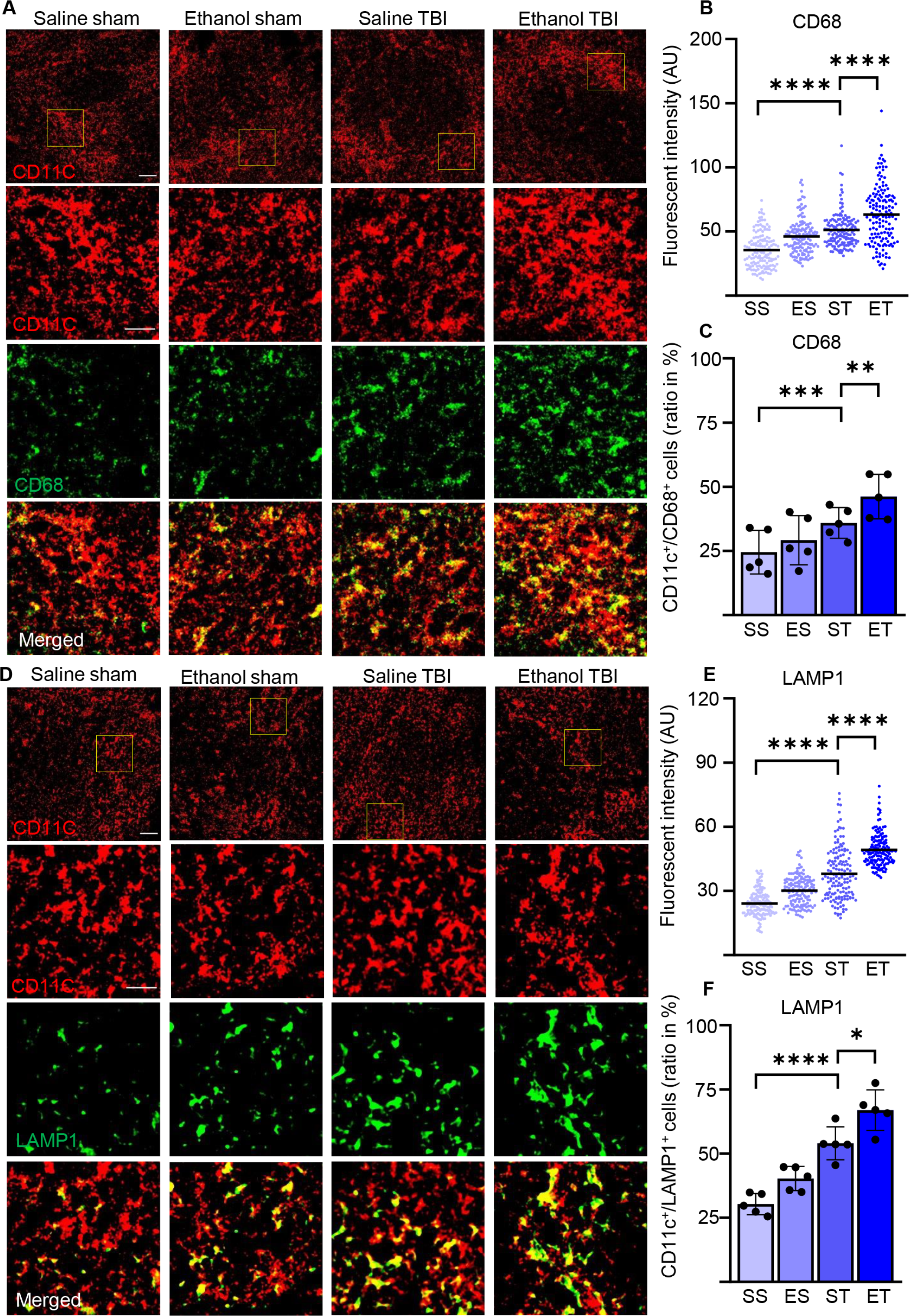
Lysosomal activity in DCs increases by TBI and is further enhanced by EI. Immunofluorescence staining of CD68 and LAMP1 with DC marker CD11c on thin spleen sections of saline sham (SS), ethanol sham (ES), saline TBI (ST) and ethanol TBI (ET) treated mice 3h after trauma. A-C) immunofluorescent staining of CD68 colocalized with CD11c resulted in a significant increase in fluorescent intensity upon TBI (SS vs ST; p < 0.0001), with a significant enhancement after EI (ST vs ET; p < 0.0001). The amount of CD68^high^/CD11c+ cells revealed a significant increase after TBI (SS vs ST; p = 0.0007), with a significant enhancement after EI (ST vs ET; p = 0.002). D-F) immunofluorescent staining of LAMP1 colocalized with CD11c resulted in a significant increase in fluorescent intensity upon TBI (SS vs ST; p < 0.0001), with a significant increase after EI (ST vs ET; p < 0.0001). The number of LAMP1^high^/CD11c+ cells revealed a significant increase after TBI (SS vs ST; p = 0.0007) with a further significant increase after EI (ST vs ET; p = 0.01). Data shown as scatterplots or barplots with individual data points. Group size: SS N = 5, ES N = 5, ST N = 5, ET N = 5. *: p < 0.05; **: p < 0.01; ***: p < 0.001; ****: p < 0.0001. Scale bar overview: 50 µm; scale bar insert: 20 µm.

### 3.6 TBI strongly induces immunogenic function of splenic DCs

The increased antigen presentation and lysosomal activity on CD11c+ DC, shown by the increase in MHC-II, CD68 and LAMP1, strongly suggests that TBI and EI induces a rapid maturation of splenic DC. However, functional aspects of these mature DCs after TBI remain unclear. Initial evidence suggested that immature DCs are tolerogenic to T cells and mature DCs increase T cell response and immunity [77, 78], however other evidence points towards mature DCs with tolerogenic function [79, 80]. To further explore the immunostimulatory phenotype of splenic DC in trauma, we assessed the expression levels of TNFα in CD11c+ DCs. In fact, TNFα is upregulated in DC by the interaction with antigens and by the stimulation of TLRs and it is a major inducer of T-cell responses [81–83]. We also considered the expression of the beta-2 adrenergic receptor, a marker associated with tolerogenic DC [84–86], under the hypothesis that expression of beta-2 adrenergic receptor and TNFα should be inversely correlated. Fluorescent single mRNA in situ hybridization was performed on spleen sections for beta-2 adrenergic receptor (ADRB2) and TNFα (Fig. 7A). Density analysis of single mRNA TNFα in CD11c+ DCs revealed a significant difference within treatment groups (Two-way ANOVA, F_(3,267)_ = 79.5; p < 0.0001), due to a strong increase after TBI (Tukey corrected, SS vs ST; 13±6 vs 22±8; p < 0.0001; Fig. 7B), with a further increase after EI (ST vs ET; 22±8 vs 29±10; p < 0.0001; Fig. 7B). Furthermore, when assessing ADRB2 in CD11c+ DCs, density analysis revealed a significant difference within treatment groups (Two-way ANOVA, F_(3,267)_ = 18.99; p < 0.0001), due to a significant decrease after TBI (Tukey corrected, SS vs ST; 14±5 vs 10±4; p < 0.0001; Fig. 7C). Interestingly, EI showed a strong increase in ADRB2 compared to TBI (ST vs ET; 10±4 vs 16±7; p < 0.0001; Fig. 7C). This data suggests, that TBI induces an increase in immunogenic function of splenic DCs, shown by the increased TNFα expression and decreased ADRB2 expression. Upon EI the immunogenic function is further enhanced shown by the increased TNFα expression, however the increase of ADRB2 suggests a simultaneous high sensitivity to adrenergic dampening of inflammation after EI.

**Figure 7:**
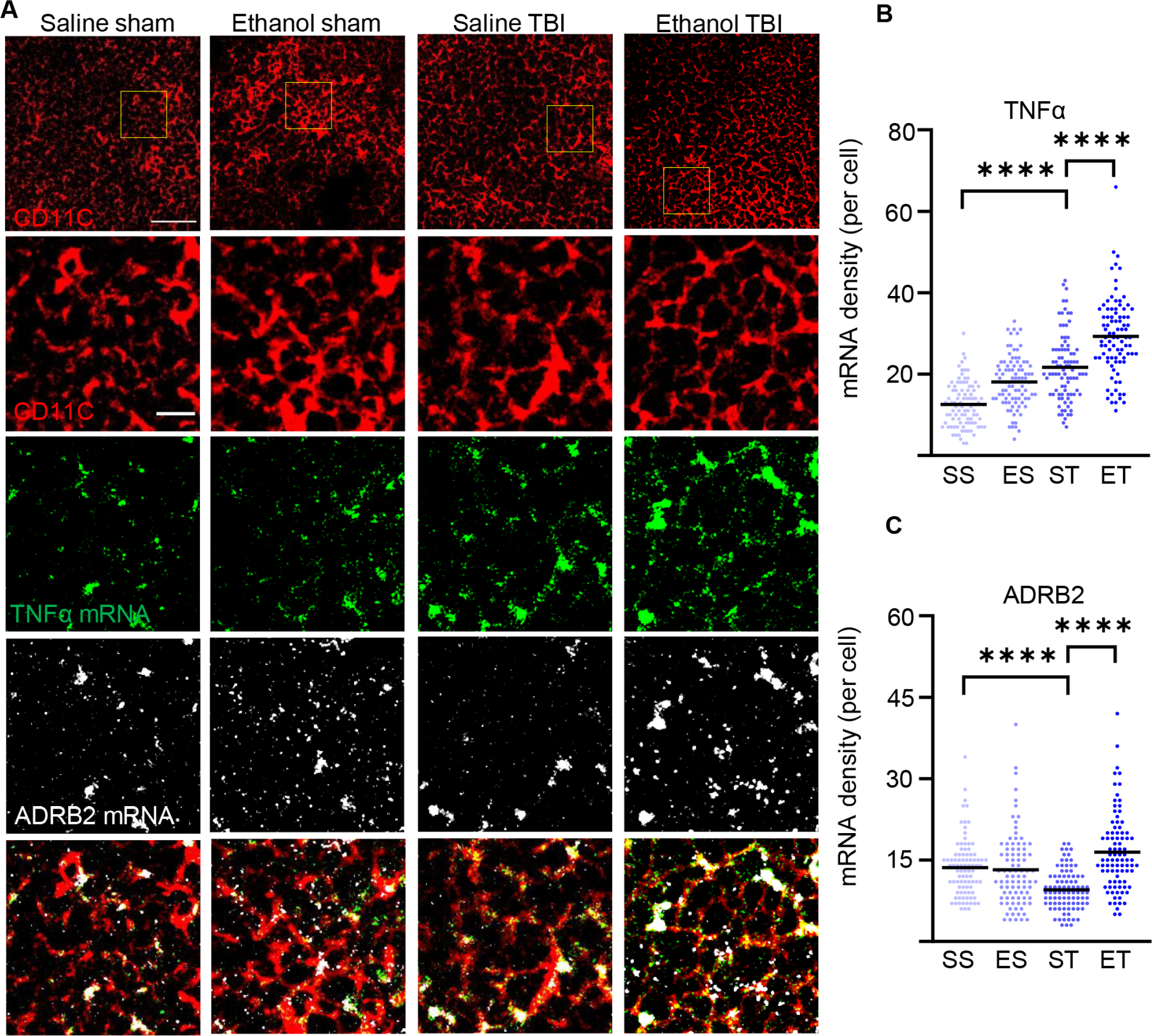
TBI enhances TNFα expression in splenic DCs and EI shows high sensitivity to adrenergic dampening. Fluorescent in situ mRNA hybridization of TNFα and beta-2 adrenergic receptor (ADRB2) with co- staining of DC marker CD11c on thin spleen sections of saline sham (SS), ethanol sham (ES), saline TBI (ST) and ethanol TBI (ET) treated mice 3h after trauma. A-B) Fluorescent in situ hybridization of TNFα resulted in a significant increase in mRNA density upon TBI (SS vs ST; p < 0.0001), with a significant enhancement after EI (ST vs ET; p < 0.0001). A-C) Fluorescent in situ hybridization of ADRB2 resulted in a significant decrease in mRNA density upon TBI (SS vs ST; p < 0.0001), however EI shows a significant increase (ST vs ET; p < 0.0001). Data shown as scatterplots. Group size: SS N = 5, ES N = 5, ST N = 5, ET N = 5. *: p < 0.05; **: p < 0.01; ***: p < 0.001; ****: p < 0.0001. Scale bar overview: 50 µm; scale bar insert: 10 µm.

## 4. Discussion

Our data show that shortly (3h) after TBI, splenic DCs undergo a maturation process that involves FLT3/FLT3L signaling, enhanced protein synthesis, increased phagocytotic and lysosome activity as well as upregulated expression of MHC-II and finally increased inflammatory properties, shown by TNFα expression. Most notably, in case of concomitant high-dose EI, the maturation process is enhanced, with increased expression of FLT3L and larger fractions of CD11c+ cells displaying elevated protein synthesis and signs of immune function activation. However, with a simultaneous increased ARB2 expression. Thus, not only the TBI sets in motion events influencing an important compartment of the systemic immune response, although concomitant EI is capable of substantially amplifying these cascades, it simultaneously shows a rapid autonomic innervation.

Maturation of DCs is conceptualized as the phenotypic change from a state characterized by high endocytic capacity, low expression of co-stimulatory molecules and MHC-II and weak induction of T-cells responses (immature dendritic cells) to a state of downregulated phagocytosis, high expression of MHC-I and MHC-II and effective stimulation of naive T cells [74]. Furthermore, maturation of DCs also involves a substantial remodeling of their metabolism, with increased mTOR-dependent protein synthesis [68, 70] and increased use of glycolytic pathways [87]. DC maturation is also associated with a significant modification in the protein degradation flux, with upregulation of lysosomal markers such as LAMP3 [88] but reduced autophagy [89]. Furthermore, the maturation process is intertwined with the upregulation of cytokine secretion such as TNF-alpha [81] and IL-12 [90]. Therefore, with the demonstration of increased FLT3 phosphorylation, upregulation of protein synthesis markers, increased expression of MHC-II and of lysosomal proteins LAMP1 and CD68, we believe to provide substantial evidence to state that TBI induces a quick upregulation of the splenic DC maturation process, from steady-state cells to effective APC.

What drives such an induction of maturation? The maturation process of DCs is set in motion, among others, by PAMPs or DAMPs [72, 73], i.e. proteins released either by bacterial or viral pathogens, or by damaged tissue of the body. Indeed, levels of brain tissue proteins and damage markers are already elevated at 3h in the serum of mice subject to TBI (including GFAP, NSE, S100B and NFL; [37, 91]. In particular, HMGB1, an alarmin located in the nucleus of neurons and glial cells and released upon brain tissue disruption [92, 93], is highly and rapidly elevated in serum after TBI [94, 95]. HMGB1 has been found to contribute not only to local neuroinflammation upon neurotrauma [96, 97] but it is also induced in non-cerebral tissues post TBI and contributes to the subsequent systemic inflammation following TBI [10]. Notably, HMGB1 is also a major inducer of DC maturation, an effect that appears to be relevant in the context of lung injury (through mTOR signaling; [98]) and in liver injury [99]. Nevertheless, recent evidence has demonstrated the strong involvement of the autonomic system in controlling splenic responses through adrenergic and cholinergic inputs [20,22,45,100,101]. Moreover, the adrenergic activation of DCs is rather associated with limited expression of MHC-II, CD86 but strongly increases the secretion of IL-10 [102]. Likewise, adrenergic stimulation of DCs substantially decreases the release of IL-12 and, in turn, suppresses the secretion of IFN-γ by Th1 lymphocytes [85]. Thus, a role for the autonomic innervation in contributing to TBI-induced splenic DC maturation may take place along with the effect of systemic, blood-borne cytokines.

What are the possible consequences of the TBI-induced maturation of splenic DCs on local and systemic immune activation following the brain injury? The net effect of the activation of the brain-spleen axis in TBI (either by circulating cytokines or by the dys-autonomia associated with TBI) seems to be detrimental, since immediate splenectomy results in an improved survival, reduced brain and systemic cytokine response, ameliorated brain oedema and preservation of cognitive abilities in experimental rat TBI [4, 103]. These effects were correlated with the decrease in NF-kB activation at the injury site [104]. Similar beneficial effects of splenectomy have been reported in the context of spinal cord injury (SCI) [105] and stroke [20]. However, the actual contribution of DC maturation may not necessarily be detrimental and may strongly depend on their activation status (immature, semi-mature or mature; [74]). It was shown that DCs pulsed with myelin-basic protein could actually drive a protective T-cell response in SCI [106]; on the other hand, autoimmune responses following TBI are associated with detrimental outcomes [107]. Interestingly, DC maturation is actually impaired in SCI patients [108]. Thus, our findings suggest that TBI causes a rapid recruitment of DCs in the spleen. Given the central role of these cells as APC, they may be substantial contributors to the systemic imbalance of immune functions following TBI.

What is the contribution of EI-driven enhanced DC maturation upon TBI? EI has been reported to be dose-dependently associated with a reduced inflammatory response (with decreased cellular inflammation and altered cytokine pattern) at the injury site [25–27] in murine TBI models. Importantly, EI also rapidly (already at 3h) dampens the systemic inflammatory response triggered by TBI. In particular, in murineTBI, levels of HMGB1 and IL-6 are decreased in the liver of mice subject to EI/TBI compared to TBI alone (but IL-1β is actually upregulated;[10]) In contrast, in the lungs EI decreases the levels of HMGB1, IL-6, IL-1β and TNF-α, while moderately increasing IL-10 [10]. In line with this evidence, EI is associated with reduced systemic IL-6 levels and less pronounced leukocytosis in human TBI patients [109] and in patients after major traumas (including TBI;[110]), as well as increased levels of IL-10 [111]. Our findings suggest that EI results in an increased number of splenic DCs undergoing a maturation process, driven by increased FLT3 phosphorylation and demonstrated by the larger fraction of CD11c+ cells displaying upregulated protein synthesis (pS6) and lysosomal markers (LAMP-1, CD68) as well as the induction of high levels of TNFα mRNA. It must be stressed that our model takes into consideration an acute consumption of a high dose of ethanol (“binge”), and therefore our findings are not directly comparable with the reports of reduced DC ability to stimulate T cells upon chronic ethanol exposure [112, 113]. Nevertheless, the combination of in vitro and in vivo data about the effects of EI on trauma-associated inflammation suggests an overall immunosuppressive effect of EI, in agreement to the impact of chronic ethanol on DC function. If this extrapolation is sound, then the enhanced maturation of splenic DCs seen in EI/TBI samples may result in a DC phenotype with inflammation-resolution properties. This hypothesis is supported by the observed upregulation of the beta-2 adrenergic receptor in DC. Nevertheless, the combination of increased TNFα and ADRB2 mRNA (not previously described), may correspond to a peculiar state of activation characterized by high immune-stimulatory properties and, at the same time, quick dampening from autonomic innervation. Thus, the impact of EI-driven expansion of DC maturation may ultimately contribute to the reduced systemic immune reactivity seen in TBI upon ethanol intoxication.

The present work is not without limitations. First, the ultimate evaluation of the DC function would require an active immunization protocol in vivo or an ex-vivo naive T-cell stimulation assay, which was beyond the technical scope of the present project. Second, the use of CD11c+ marker does not distinguish the subpopulation of myeloid DC or the plasmacytoid DC; we have nevertheless maintained a consistent selection of the region of interest in correspondence of the marginal zone [114].

In conclusion, our findings show that induction of maturation markers in splenic DC takes place rapidly after TBI and is highly correlated with the phosphorylation of FLT3; we further demonstrate that concomitant EI amplifies the maturation process of splenic DC post TBI. Thus, our findings identify the DC as a new player in the immunomodulation occurring upon EI in TBI.

## Acknowledgement

We thank all the members of the CRC 1149 for their scientific input and discussion. We would like to thank prof. Frank Kirchoff for the use of the confocal facility and prof. Anita Ignatius for the access to the histology facility. Technical support by Thomas Lenk was highly appreciated.

## Authorścontribution statement

FR and FoH conceived and designed the project. JZ, ZL and SL performed the analysis of the spleen tissue. AC performed the TBI procedures. AL, TB, MHL, FR and FoH contributed to the analysis and the interpretation of the data. JZ, FR, MHL and FoH wrote the first draft of the manuscript. FR, FoH, JZ, ZL, SL, AL, TB, MHL contributed to the final version of the manuscript.

## Figure legends

**Supplementary Figure 1: CD11c+ DC density is unaffected by TBI and EI. FLT3 phosphorylation remains unaltered in CD45+/CD11c- cells.**

Immunofluorescence staining of thin spleen sections of saline-sham (SS), ethanol-sham (ES), saline-TBI (ST) and ethanol-TBI (ET). A) Immunofluorescence staining of CD11c in control groups reveals inhomogeneous distribution in thin spleen sections. With a high localisation around the follicles. B-C) Density of CD11c+ cells remains unaltered 3h after TBI and EI (p = 0.45). D-E) immunofluorescence staining of CD45, CD11c and pFLT3 reveals unaltered phosphorylation levels of FLT3 in CD45+/CD11c- cells (p = 0.50). Data shown as barplots with individual data points. Group size: SS N = 5, ES N = 5, ST N = 5, ET N = 5. *: p < 0.05; **: p < 0.01; ***: p < 0.001; ****: p < 0.0001. Scale bar overview A: 200 µm; scale bar insert A: 50 µm; scale bar overview B and D: 50 µm; scale bar insert D: 20 µm.

**Supplementary Table 1: Primer sequences used for gene expression analysis (RT-qPCR).**

**Supplementary Table 2: Primary and secondary antibodies used for immunostaining (IF).**

