## Supplementary figures and images for "Fast maturation of splenic dendritic cells upon TBI is associated with FLT3/FLT3L signaling"

### supplementary figure 1

Zhang et al., Supplementary Figure 1.pdf
2050 x 2714

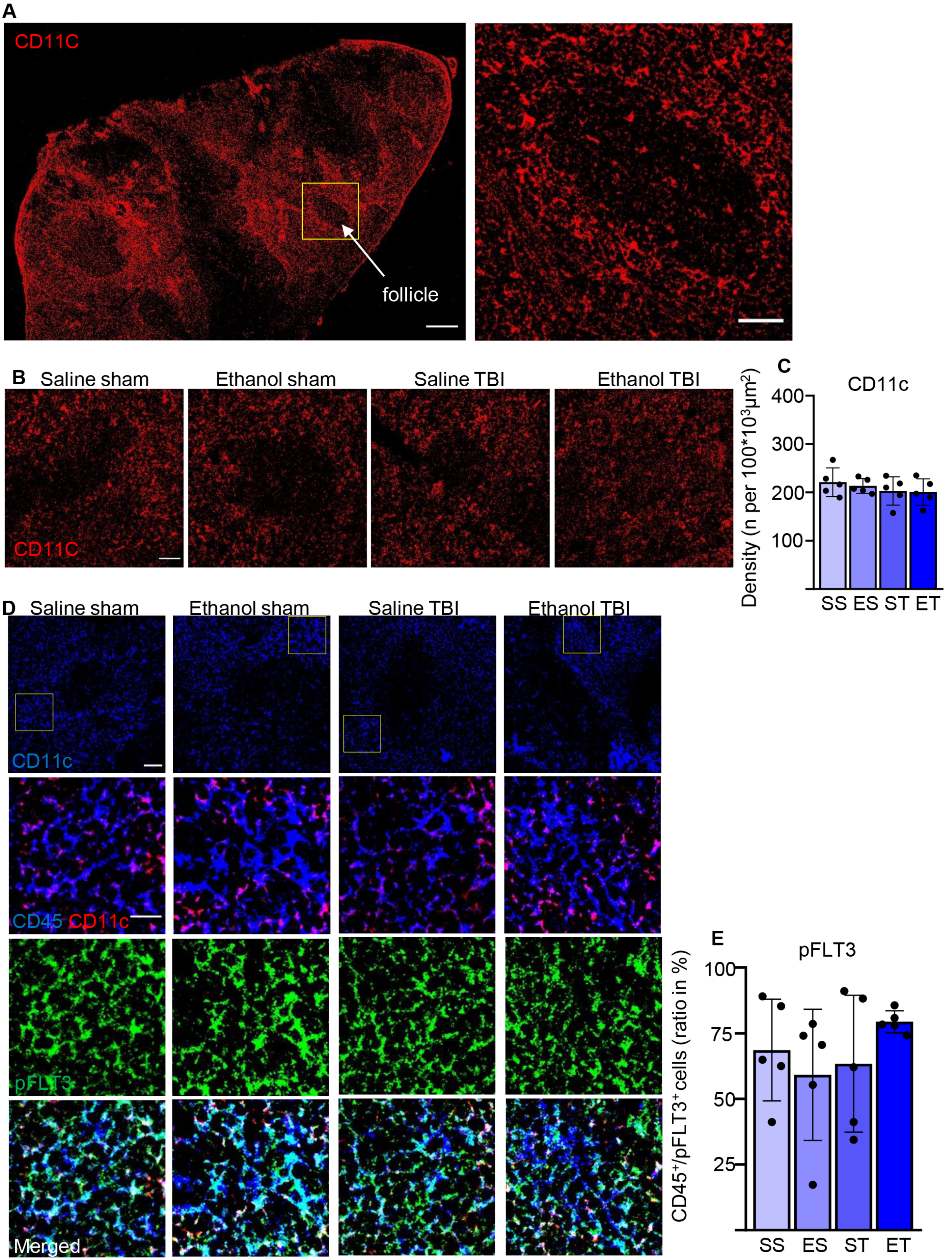
