## supplementary table 1 for "Fast maturation of splenic dendritic cells upon TBI is associated with FLT3/FLT3L signaling"

**Table 1: List of primer sequences.**

| **Gene** | **Sequence** |
| --- | --- |
| IL-13 | forward: 5’ -acc cag agg ata ttg cat gg - 3’  reverse: 5’ - tgg gct act tcg att ttg gt - 3’ |
| IL-4 | forward: 5’ -gag aga tca tcg gca ttt tga - 3’  reverse: 5’ -agc cct aca gac gag ctc ac- 3’ |
| IL-10 | forward: 5‘ -agg cgc tgt cat cga ttt ctc- 3‘  reverse: 5‘ - gcc ttg tag aca cct tgg tct t- 3‘ |
| IL-19 | forward: 5’ -gga gca ata gac agc ccc tc- 3’  reverse: 5’ -tgc act aca gca cac cac aa- 3’ |
| IL-12 | forward: 5’ -cct gtg cct tgg tag cat ct- 3’  reverse: 5’ –ctg aag tgc tgc gtt gat gg- 3’ |
| IL-17 | forward: 5’ -gga cag ccc ttc ttt gtc tg- 3’  reverse: 5’ -tgc ttt tta tat ttc att acg tgg tt- 3’ |
| IL-23a | forward: 5’ -gag caa ctt cac acc tcc cta- 3’  reverse: 5’ -tag aac tca ggc tgg gca tc- 3’ |
| IFN-y | forward: 5’ -ttc aag act tca aag agt ctg agg- 3’  reverse: 5’ -tct gga gga act ggc aaa ag- 3’ |
| FLT3L | forward: 5’ -gct ctg aag ccc tgt atc gg- 3’  reverse: 5’ -tgg ctt cta ggg cta tgg ga- 3’ |
| FLT3 | forward: 5’ -tgt cag taa tga ttc ttg aga ccg- 3’  reverse: 5’ -ctt ctg ggg atc ctc gca c- 3’ |
| CX3CL1 | forward: 5’ -gca agt ttg aga agc ggg tg- 3’  reverse: 5’ -att cag gct ttg tca ggg ca- 3’ |
| CX3CR1 | forward: 5’ -tca ccg tca tca gca tcg ac- 3’  reverse: 5’ -cgc cca gac taa tgg tga ca- 3’ |
| CCL2 | forward: 5’ -caa atg tgc ttg cct gac cc- 3’  reverse: 5’ -tcc tca ttt ggg gcc tga ac- 3’ |
| CCL3 | forward: 5’ -tgc cct tgc tgt tct tct ct- 3’  reverse: 5’ -gtg gaa tct tcc ggc tgt ag- 3’ |
| CCL4 | forward: 5’ -ccc agc tct gtg caa acc ta- 3’  reverse: 5’ -cca ttg gtg ctg aga acc ct- 3’ |
| CCL12 | forward: 5’ -aat cac aag cag cca gtg tcc -3'  reverse: 5’ -gtc agc aca gat ctc ctt atc cag t -3' |
| CCL1 | forward: 5’ -tgc cgt gtg gat aca gga tg- 3’  reverse: 5’ -aca gga gga gcc cat ctt tc- 3’ |
| CCL22 | forward: 5’ -gtg gaa gac agt atc tgc tgc c- 3’  reverse: 5’ -agg ctt gcg gca gga ttt tga g- 3’ |
| CXCL10 | forward: 5` -gct gcc gtc att ttc tgc- 3´  reverse: 5´-tct cac tgg ccc gtc atc- 3´ |
| CCL20 | forward: 5’ – gtg ggt ttc aca aga cag atg gc - 3’  reverse: 5’ -cca gtt ctg ctt tgg atc agc g- 3’ |
| CCL6 | forward: 5’ - tga gaa act cca aga ctg cca t  reverse: 5’ -tgg agg gtt ata gcg acg atc t |
| CCL24 | forward: 5’ – cag cct tct aaa ggg gcc aa - 3’  reverse: 5’ -cta aac ctc ggt gct att gcc- 3’ |
| CXCL3 | forward: 5‘ -gaa agg agg aag ccc ctc ac- 3‘  reverse: f‘ -aca cat cca gac acc gtt gg- 3‘ |
| CXCL16 | forward: 5‘ -gca ggg tac ttt gga tca cat cc- 3‘  reverse: 5‘ -agt tca cgg acc cac tgg tct t- 3‘ |
