## supplementary table 2 for "Fast maturation of splenic dendritic cells upon TBI is associated with FLT3/FLT3L signaling"

**Table 2: List of antibodies.**

| **Antigen (primary antibody)** | **Catalogue number** | **Company** | **Dilution** |
| --- | --- | --- | --- |
| Rabbit anti CD11c | 375 003 | Synaptic Systems | 1:500 |
| Mouse anti CD11c | ab11029 | Abcam | 1:100 |
| Rat anti CD45 | 550539 | BD Bioscience | 1:50 |
| Rabbit anti FLT3 (phospho) | bs-3149R | Bioss | 1:100 |
| Rabbit anti BTK (phospho) | NBP1-78295 | Novus | 1:200 |
| Rabbit anti S6-RP (phospho) | 2211S | CST | 1:200 |
| Rabbit anti eIF2A (phospho) | 3597S | CST | 1:50 |
| Mouse anti MHC-II | 12-5321-82 | Invitrogen | 1:200 |
| Mouse anti CD68 | Ab31630 | abcam | 1:200 |
| Rat anti LAMP1 | 1D4B | DSHB | 1:30 |
| **Antigen (secondary antibody)** | **Catalogue number** | **Company** | **Dilution** |
| Donkey anti mouse 488 | A21202 | Invitrogen | 1:500 |
| Donkey anti mouse 568 | A10037 | Invitrogen | 1:500 |
| Donkey anti mouse 647 | A31571 | Invitrogen | 1:500 |
| Donkey anti rabbit 488 | A21206 | Invitrogen | 1:500 |
| Donkey anti rabbit 568 | A10042 | Invitrogen | 1:500 |
| Donkey anti rabbit 647 | A31573 | Invitrogen | 1:500 |
| Chicken anti rat 568 | A21472 | Invitrogen | 1:500 |
